# Altered Bacteria-Fungi Inter-Kingdom Network in Gut of Ankylosing Spondylitis Patients

**DOI:** 10.1101/396812

**Authors:** Ming Li, Bingbing Dai, Yawei Tang, Lei Lei, Ningning Li, Chang Liu, Teng Ge, Lilong Zhang, Yao Xu, Yuqi Hu, Pengfei Li, Yan Zhang, Jieli Yuan, Xia Li

## Abstract

Intestinal bacterial dysbiosis has been increasingly linked to Ankylosing Spondylitis (AS), which is a prototypic and best studied subtype of Spondyloarthritis (SpA). Fungi and bacteria coexist in human gut and interact with each other, although they have been shown to contribute actively to health or diseases, no studies have investigated whether fungal microbiota in AS patients is perturbed. In this study, fecal samples of 22 AS patients, with clinical and radiographic assessments, and 16 healthy controls (HCs) were collected to systematically characterize the gut microbiota and mycobiota in AS patients by 16S rDNA and ITS2-based DNA sequencing. The relationships between therapeutic regimens, disease activity, radiographic damage of AS and gut micro/mycobiome were investigated. Our results showed a distinct mycobiota pattern in AS in addition to microbiota dysbiosis. The gut mycobiome of AS patients was characterized by higher taxonomic levels of *Ascomycota*, especially the class of *Dothideomycetes*, and decreased abundance of *Basidiomycota*, which was mainly contributed by the decease of *Agaricales*. Compared to HCs, changing of the ITS2/16S biodiversity ratio, and bacteria-fungi interkingdom network were observed in AS patients. Alteration of gut mycobiota was associated with different therapeutic regimens, disease activity, as well as different degrees of radiographic damage. Moreover, we unraveled a disease-specific interkingdom network alteration in AS. Finally, we also identified some trends suggesting that different therapeutic regimens may induce changing of both bacterial and fungal microbiota in AS.

**IMPORTANCE:** Human gut is colonized by diverse fungi (mycobiome), and they have long been suspected in the pathogenesis of Spondyloarthritis (SpA). Our study unraveled a disease-specific interkingdom network alteration in AS, suggesting that fungi, or the interkingdom interactions between bacteria and fungi, may play an essential role in AS development. However, limited by sample size and indeep mechanism studies, further large scale investigations on the characterization of gut mycobiome in AS patients are needed to form a foundation for research into the relationship between mycobiota dysbiosis and AS development.

## INTRODUCTION

Spondyloarthritis (SpA) is a group of several related but phenotypically distinct disorders: psoriatic arthritis (PsA), arthritis related to inflammatory bowel disease (IBD), reactive arthritis, a subgroup of juvenile idiopathic arthritis, and ankylosing spondylitis (AS) (1). The exact pathogenesis of SpA remains unknown (2); however, altered immune responses towards gut microbiota under the influence of genetic and environmental factors have been shown in autoimmune diseases related to SpA (3-6).

Among the related disorders, AS is the prototypic and best studied subtype of SpA. Up to 70% of AS patients have subclinical gut inflammation and 5-10% of these patients have more severe intestinal inflammation that progresses to clinically defined IBD (7). As intestinal dysbiosis has been increasingly linked to IBD in recent years (8-10), it is reasonable to speculate a close link between gut microbiota and AS development (3,11). Previous works have shown that the patients and transgenic rat model of AS had increased immunoglobulins G (IgG) or pro-inflammatory cytokines in response to bacterial products such as outer membrane protein and lipopolysaccharide (LPS) (12,13). A small case 16S ribosomal DNA sequencing analysis has shown dysbiosis in terminal ileum biopsy specimens of AS patients (14). A recent quantitative metagenomics study, based on deep shotgun sequencing using gut microbial DNA from 211 Chinese individuals, also proved that alterations of the gut microbiome were associated with development of AS (15). Alterations of gut microbial genera, such as *Bacteroides* (16), *Prevotella* (17), *Bifidobacterium* and *Lachnospiraceae* subgroups, *etc*. (18) in IBD were highly in accordance with the patterns that were observed in AS patients.

Besides bacterial dysbiosis, a distinct alteration of fungal microbiota (mycobiota) was also identified in fecal samples of IBD patients (19). Although constituting only a small part of gut microbiome (20), mycobiota has been shown to contribute actively to health or diseases in a complex manner (21,22). Actually, fungi have long been suspected in SpA. For example, the anti-*Saccharomyces cerevisiae* antibodies (ASCA) were found to be associated with intestinal inflammation in SpA (23). β-1,3-glucan, a fungal product, had been shown to trigger SpA in BALB/c ZAP-70W163C–mutant (SKG) mice (24), and this response was mediated by interleukin-23 (IL-23)-provoked local mucosal dysregulation and cytokines driving SpA syndrome (25). Dectin-1, the C-type lectin-like pattern recognition receptor of β-1,3-glucan, and downstream gene Caspase recruitment domain-containing protein 9 (CARD9) are the common candidates for genetic studies in AS, PsA and Crohn’s disease (26,27). However, to our knowledge, no studies have investigated whether fungal microbiota in AS patients is perturbed.

In this study, we characterized both microbial and fungal microbiota in fecal samples of AS patients using high through-out sequencing, and analyzed the correlation between bacterial and fungal microbiota. We also compared the gut microbiome of AS patients receiving different therapeutic regimens, or with different disease activities. Data in our study represent a first systematic analysis of microbiome in AS patients, and provide a rationale to support the role of mycobiota dysbiosis in AS pathogenesis.

## RESULTS

### Participant characteristics

We included a total of 71 individuals in the current analysis, composed of 30 healthy controls (HCs), 41 cases with AS (Supplementary Fig. S1). Of the AS cases, 19.51 % (n=8) were newly diagnosed as AS, 80.49 % (n =33) were patients that had different years of disease duration, and were treated by biological agents (BLs) or NSAIDs. Due to medical histories of other diseases, and/or use of antibiotics, probiotics, prebiotics or synbiotics before fecal samples collection, 19 patients were excluded. The included 22 AS patients are all males with an average age of 34.86 years old. 14 HCs were excluded for age and gender matching, the remained HCs are all males with an average age of 34.35 (No age difference between AS and HC group, p>0.05). As expected, the majority of these patients were detected as HLA-B27 positive (>85 %) and with axial involvement (>94 %). The disease activity parameters including CRP, ESR, and BASDAI were summarized in Table 1. The radiographic assessments showed that 22.73 %, 45.45 %, and 31.82 % of the patients have II, III, and IV levels of structural damage in their spine, respectively. During follow-up, 9 of the patients were at some time exposed to NSAIDs and 8 to BLs such as TNF inhibitors.

**TABLE 1.** Baseline demographic, clinical and radiographic characteristics of the AS patients.

| Characteristic | AS patients (n=22) |
| --- | --- |
| Age, years, mean (range) | 34.86 (15-58) |
| Male, % | 100 |
| Disease duration, years, mean (range) | 9.60 (0.17-40) |
| HLA-B27 positive, % | 85.71 |
| **Disease activity parameter** |  |
| CRP, mg/dl, mean (range) | 14.52 (0.1-67) |
| ESR, mm/h, mean (range) | 27.53 (2-67) |
| Axial involvement, % | 94.44 |
| BASDAI, mean (range) | 5.05 (2.1-9.4) |
| **Imaging classification** |  |
| I, % | 0 |
| II, % | 22.73 |
| III, % | 45.45 |
| IV, % | 31.82 |
| **Medication use** |  |
| NSAIDs, % | 40.91 |
| Biological agent, % | 36.36 |
| Treatment naïve, % | 22.73 |

### Altered bacterial microbiota in AS patients

We first analyzed the bacterial fraction of the microbiota using high-throughput sequencing of the bacterial 16S ribosomal RNA gene. Compared with HCs, the observed species and alpha diversity (assessed using Shannon and Simpson index) of gut microbiota in AS patients was relatively increased, while there were no statistical differences among all indexes (Fig.1A, B, C, all p>0.05). The analysis based on weighted unifrac showed a statistically significant increase of beta diversity in AS group compared with HC group (Fig. 1D, p=0.0022), although the NMDS analysis did not exhibit an obvious separation between AS samples and those of HC group (Fig. 1E).

**FIG 1:**
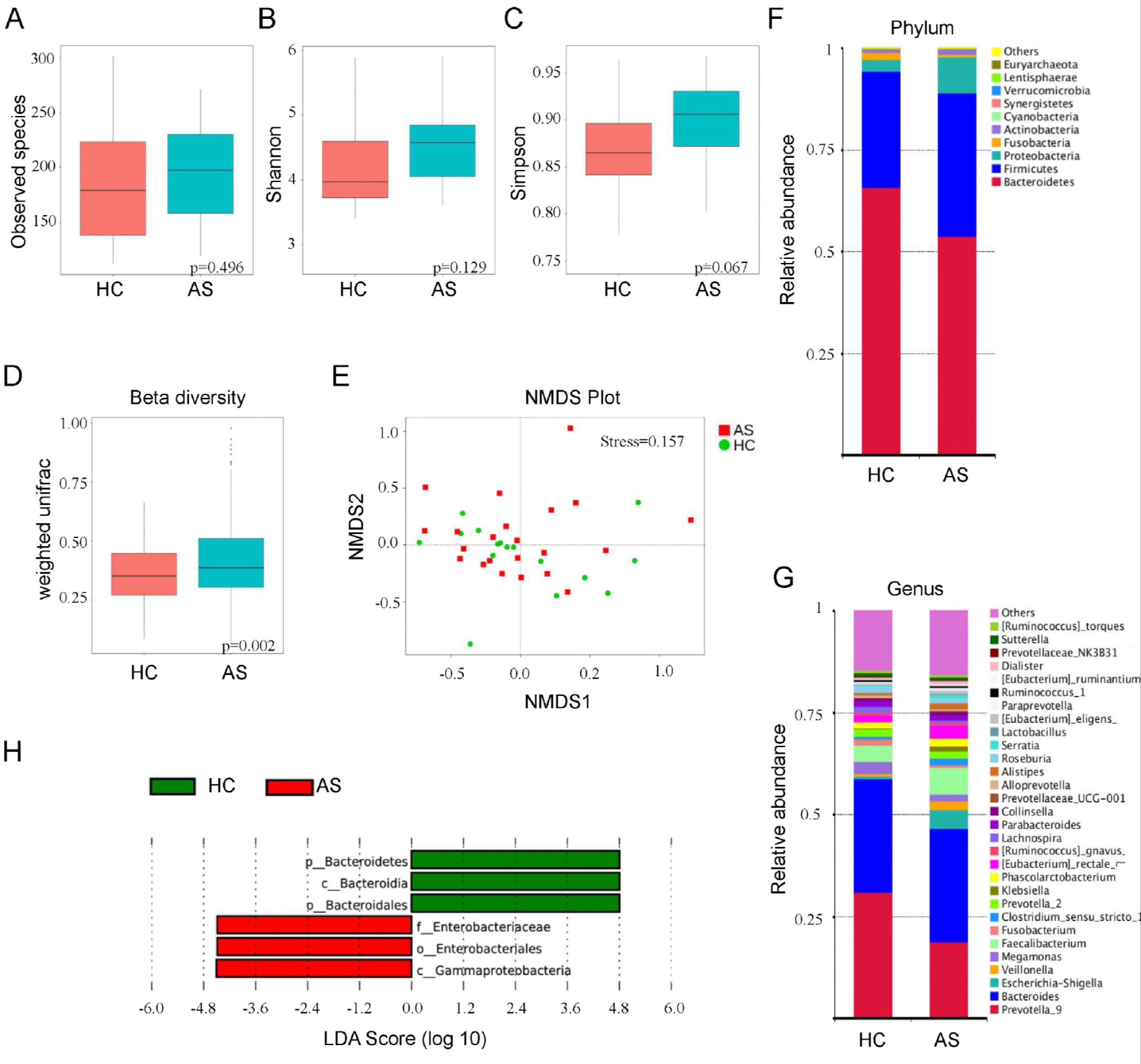
Altered bacterial microbiota biodiversity and composition in AS. (A,B and C) Observed species, Shannon, Simpson index describing the alpha diversity of the bacterial microbiota in two groups. (D) Beta diversity. (E) NMDS analaysis (F and G) Global composition of bacterial microbiota at the phyla and genus levels. (H) Taxa differentiating AS from HC.

The analysis of phylotypes indicated that *Bacteroidetes*, *Firmicutes*, *Proteobacteria*, *Fusobacteria*, and *Actinobacteria* were the dominant taxa in both the AS patients and healthy controls (Fig. 1F). At the phylum level, increased abundance of *Proteobacteria* (p=0.0399) and decreased *Bacteroidetes* (p=0.0177) were found in AS patients compared with HC group (Supplementary Fig. S2 and Table S1). We also observed a greater abundance of *Firmicutes, Actinobacteria*, and a lower abundance of *Fusobacteria*, which highly agree with the results of Wen *et al*. (15), although the t-test between groups showed insignificant (all p>0.05). Consistent with this result, enriched bacterial genera of *Escherichia*-*Shigella* (*Proteobacteria*), *Veillonella* (*Firmicutes*), *Faecalibacterium* (*Firmicutes*), *Eubacterium* rectale group (*Firmicutes*), *Streptococcus* (*Firmicutes*), *Lachnospiraceae* NK4A136 group (*Firmicutes*), and reduced patern of *Prevotella* 9 (*Bacteroidetes*), *Megamonas* (*Firmicutes*), *Fusobacterium* (*Fusobacteria*) were detected (Fig. 1G).

A LefSe analysis was further adopted to identify the bacterial groups that showed significant differences in abundance between AS and HC. As shown in Fig. 1H, the comparison between AS and HC groups revealed that the major depleted bacterial group in AS patients is the phylum of *Bacteroidetes*, especially the class of *Bacterioidia* and order of *Bacteroidales*. In contrast, *Enterobacteriales* and *Gammaproteobacteria* were significantly abundant in AS (Supplementary Fig. S3).

### Altered bacterial microbiota in AS patients receiving different therapeutic regimens

In our study, some of the fecal samples were from newly diagnosed AS patients without any medical treatment, and they were defined as the treatment naïve group (TN, n=6). The other patients were grouped by the therapeutic regimens received, including biologics (BL, n=8) and NSAID (NS, n=9) (Details in Table 1). Compared with healthy individuals, the species of gut bacteria in AS patients treated with NSAID were obviously increased, but the statistic test did not show significant difference between any of the two groups (Fig. 2A, all>0.05, Supplementary Table S2). The Shannon and Simpson index suggested a significant elevation of alpha diversity in treatment naïve patients (p=0.0412, p=0.0158, Supplementary Table S3) compared with HC group. Among the AS patients, treatment of biologics resulted in reduction of alpha diversity of gut bacteria in contrast to the TN group, especially when tested by Simpson index (p=0.0497, Supplementary Table S4). The principal coordinate analysis (PCoA) by both weighted and unweighted UniFrac showed that there was no obvious separation of groups (Fig. 2B). The variations of gut microbes in each group were also observed on phylum and genus levels (Fig. 2C, D). The *Proteobacteria* (especially the family of *Enterobacteriaceae*) and the genus of *Veillonella* were found enriched in AS patients treated with biologics when compared with the patients without treatment (TN group), in which the species of *Phascolarctobacterium faecium* was significantly more abundant (Fig. 2E). However, no bio-markers were detected in others groups of AS patients by means of LefSe analysis.

**FIG 2:**
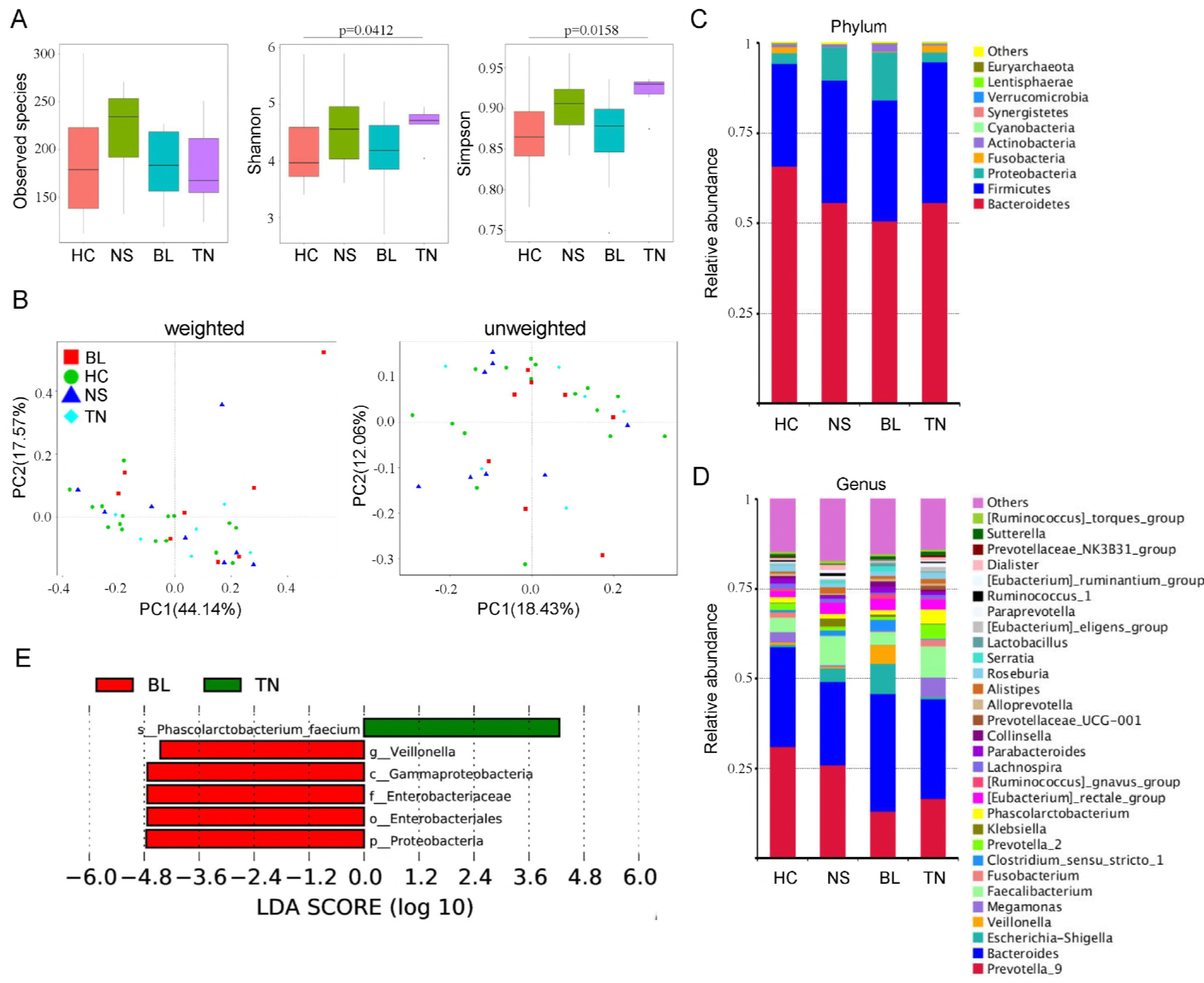
Altered bacterial microbiota biodiversity and composition in AS patients receiving different therapeutic regimens. (A) Observed species, Shannon index and Simpson index describing the alpha diversity of the bacterial microbiota in the different groups. (B) Beta diversity. Principal coordinate analysis (PCoA) of Bray–Curtis distance with each group coloured according to the different treatment methods. PC1 and PC2 represent the top two principal coordinates that captured most of the diversity. The fraction of diversity captured by the coordinate is given as a percentage. Groups were compared using Permanova method. (C and D) Global composition of bacterial microbiota at the phylum and genus levels. (E) Taxa differentiating AS-BL group from AS-TN group.

### Altered mycobiota in AS patients

By sequencing of ITS2, we assessed the composition of fungal microbiota in our population. The results showed that, there were 354 OTUs unique to HC, 265 OTUs unique to AS, while 349 were shared between two groups (Fig. 3A). Different from the results found with the bacterial microbiota, the alpha diversity of intestinal fungi was significantly decreased in AS patients, as shown in Fig. 3B, the observed species and shannon index in AS group were significantly lower than in controls (All p<;0.05, Supplementary Table S5). To explore the equilibrium between bacteria and fungi diversity in the gut, we determined the fungi-to-bacteria species ratio. This ratio was significantly decreased in AS samples (p=0.027, Fig. 3C). The PCoA showed that AS samples grouped separately from HC, indicating that changing of fungal communities might be one of the factors influencing the disease (Fig. 3D). A detailed comparison of relative abundance of fungi between HC and AS (Fig. 3E) showed that, the phyla of *Ascomycota* and *Basidiomycota* were dominated in both groups, and there was an obvious changing in proportion of these two phyla in AS patients. Among the most dominant genera, *Alternaria, Saccharomyces* and *Candida* were increased in AS patients, in contrast to decrease in other genera (Fig. 3F). The comparison between AS and HC by LefSe revealed that the higher taxonomic levels of *Ascomycota*, especially the class of *Dothideomycetes* in this phylum, were significantly more abundant in AS patients (Fig. 3G), except the family of *Xylariaceae* (which belong to *Ascomycota*), while the phylum of *Basidiomycota* was dominant in HC, that may mainly contribute by the abundance of *Agaricales*.

**FIG 3:**
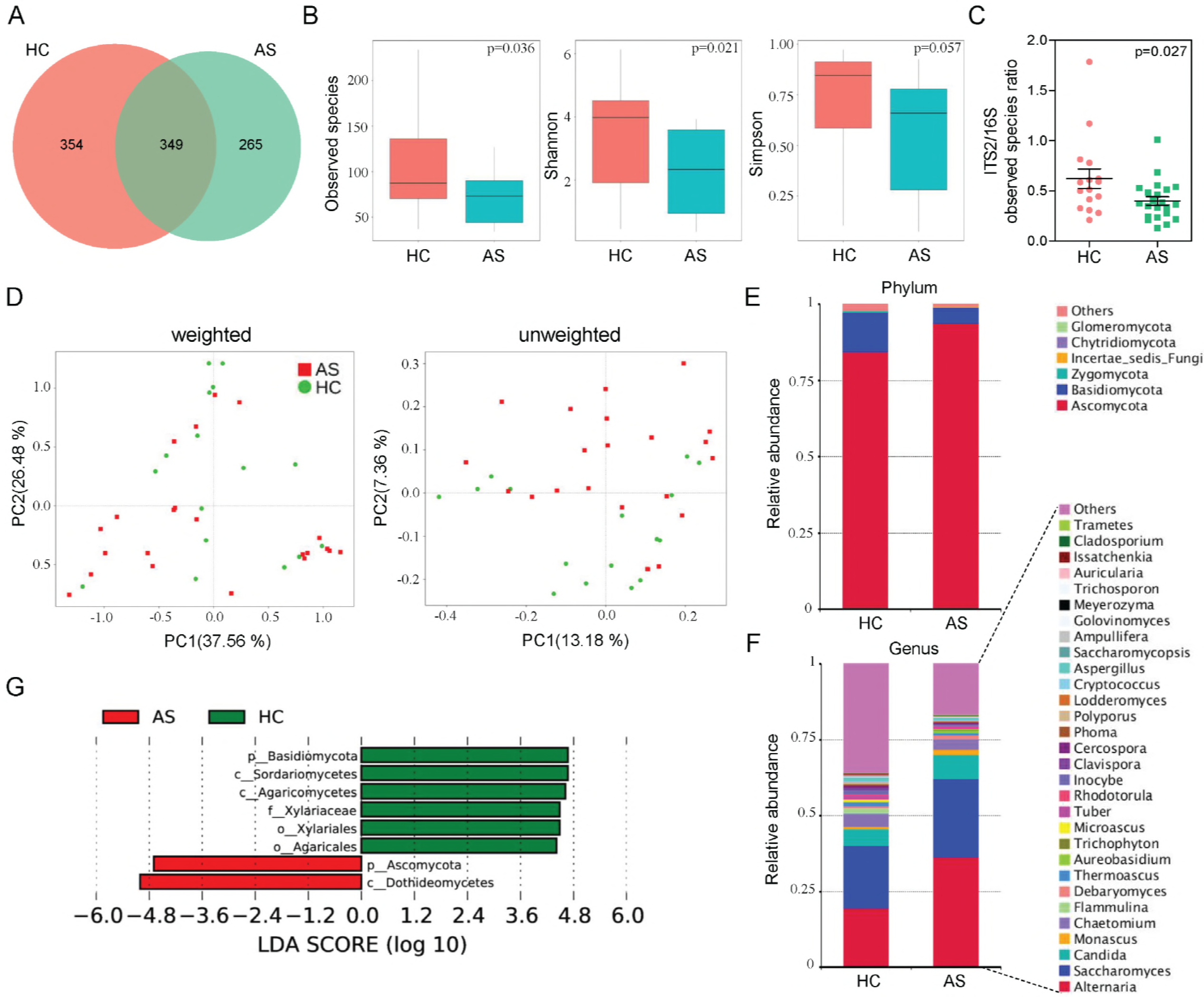
Altered fungal microbiota biodiversity and composition in AS. (A)The Venn diagram depicts OTUs that were unique to HC, unique to AS or shared. (B) Observed species, Shannon, Simpson index describing the alpha diversity of the fungal microbiota in two groups. (C) ITS2/16S observed species ratio. (D) Beta diversity. PCoA of Bray–Curtis distance with each sample coloured according to the two groups. PC1 and PC2 represent the top two principal coordinates that captured most of the diversity. The fraction of diversity captured by the coordinate is given as a percentage. Groups were compared using Permanova method. (E and F) Global composition of fungal microbiota at the phyla and genus levels. (G) Taxa differentiating AS from HC samples.

### Altered mycobiota in AS patients receiving different therapeutic regimens

We further compared the gut mycobiota of AS patients grouped by different therapeutic regimens. Notably, a significantly reduced alpha diversity was observed in treatment naïve AS patients when compared with the healthy control, especially when evaluated by Shannon and Simpson index (Fig. 4A, all p<;0.05, Supplementary Table S6-8). Treatment with biologics resulted an even lower level of observed fungal species and alpha diversity compared with the untreated TN group. In contrast, the NSAID treatment did not induce a distinct change in the number of gut fungal species in AS patients. However, when evaluated by the fungi-to-bacteria diversity ratio, we observed a significant decreased pattern in AS patients treated with both NSAID (Fig. 4B, p=0.027) and biologics (p=0.046), compared with that of the HC group. The PCoA by both weighted and unweighted Unifrac showed that the gut mycobiota of BL group separately clearly from HC, NS, and TN groups, indicating that the treatment of biologics has a profound influence on the fungal communities in AS patients (Fig. 4C). The LefSe analysis revealed that the most dominant fungal microbiota differs significantly among the four groups (Fig. 4D). Notably, the fungal microbiota in treatment naïve AS patients is characterized by the dominant of *Dothideomycetes* class, which is consist with the results of Fig. 3. In BL group, the most dominant fungal microbiota was *Saccharomyces*, this genus contributed significantly to the abundance of *Ascomycota* in AS patients of BL group.

**FIG 4:**
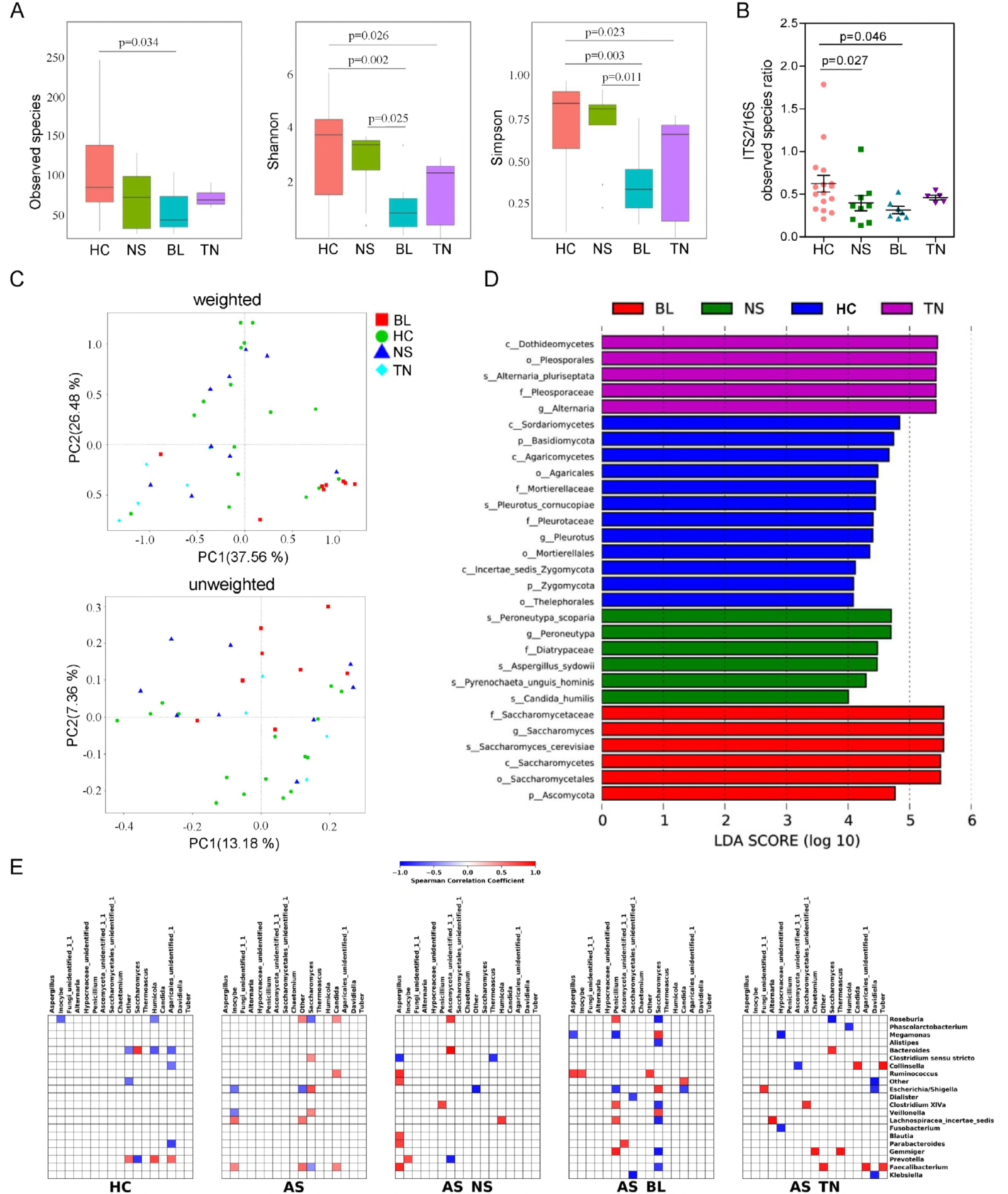
Altered mycobiota and bacteria–fungi correlation in AS patients receiving different therapeutic regimens. (A) Observed species, Shannon index,Simpson index describing the alpha diversity of the fungal microbiota in four groups studied. (B) ITS2/16S observed species ratio. (C) Beta diversity. PCoA of Bray-Curtis distance with each sample coloured according to the four groups. PC1 and PC2 represent the top two principal coordinates that captured most of the diversity. The fraction of diversity captured by the coordinate is given as a percentage. Groups were compared using Permanova method. (D) The main composition of fungal microbiota in four groups studied. (E) Specific bacteria-fungi correlation pattern in AS. Distance correlation plots of the relative abundance of fungi and bacteria genera. Statistical significance was determined for all pairwise comparisons; only significant correlations (p value <;0.05 after false discovery rate correction) are displayed. Positive values (blue squares) indicate positive correlations, and negative values (red squares) indicate inverse correlations. The shading of the square indicates the magnitude of the association; darker shades are more strongly associated than lighter shades. The sign of the correlation was determined using Spearman’s method.

### AS patients showed altered bacteria - fungi associations

In addition to composition differences, we found that the bacterial and fungal microbiota network at genus level in AS patients was notably different from that in healthy controls (Supplementary Fig. S4). Specifically, the density of bacterial network in AS patients was remarkable higher than that of the healthy individuals, while reduced network centralization and density of fungal communities were detected in these patients, which suggested an alteration of entire ecosystem in gut of AS patients. To test this hypothesis, we further investigated the bacteria-fungi correlation at the genus level according to disease phenotype. A higher spearman correlation in AS compared with HC was found (Fig. 4E). Interestingly, in AS patients, we observed a positive correlation between the abundance of *Saccharomyces* and *Clostridium sensu stricto*, *Escherichia*/*Shigella, Veillonella*, while a negative correlation between the abundance of *Saccharomyces* and *Roseburia* and *Faecalibacterium*. A positive correlation between the abundance of *Candida* and *Roseburia, Faecalibacterium* and *Ruminococcus* was also detected in AS patients, which differed from that of the HC group. Strikingly, among the AS patients, treatment of biologics and NSAID induced extensive changes in bacteria-fungi associations when compared with the untreated AS patients. Notably, many positive correlations connecting genera from *Aspergillus* to *Ruminococcus*, *Blautia*, *Parabacteroides* and *Faecalibacterium* were observed in AS patients with NSAID treatment. And there was more positive correlation between the abundance of Penicillium and *Clostridium* XIVa, *Roseburia*, *Lachnospiracea incertea sedis* and *Gemmiger* in AS patients with biologics treatment. Additionally, *Saccharomyces* followed a complicated opposite correlation with several bacterial genera in BL group. Taken together, these results suggest a complex relationship between the bacteria and fungi in the gut microbiota, and that specific alterations are present in patients receiving different therapeutic regimens.

### Altered mycobiota in AS patients was associated with disease activities and degree of radiographic damage

The canonical correspondence analysis (CCA) was used to establish the relationship between AS disease activity indexes (including BASDAI, CRP, and ESR) and the bacterial and fungal genera. As shown in Fig. 5A, the BASDAI and CRP levels were found strongly correlated to the fungal genera in treatment naïve AS patients (TN), whereas no obvious correlations were detected between bacteria genera and the disease activity indexes. We further analyzed the gut bacterial and fungal compositions of AS patients at genus level according to their stages of radiographic changes by principal component analysis (PCA, Fig. 5B). Intriguingly, a strongly separated pattern of gut mycobiota was observed in AS patients with Grade III and Grade IV stages, when compared with the healthy controls and the Grade II stage of AS patients. The elevated relative abundance of genera, such as *Saccharomyces* and *Lodderomyces*, in AS patients at Grade IV stage may contribute to the alteration of fungal community patterns (Supplementary Fig. S5).

**FIG 5:**
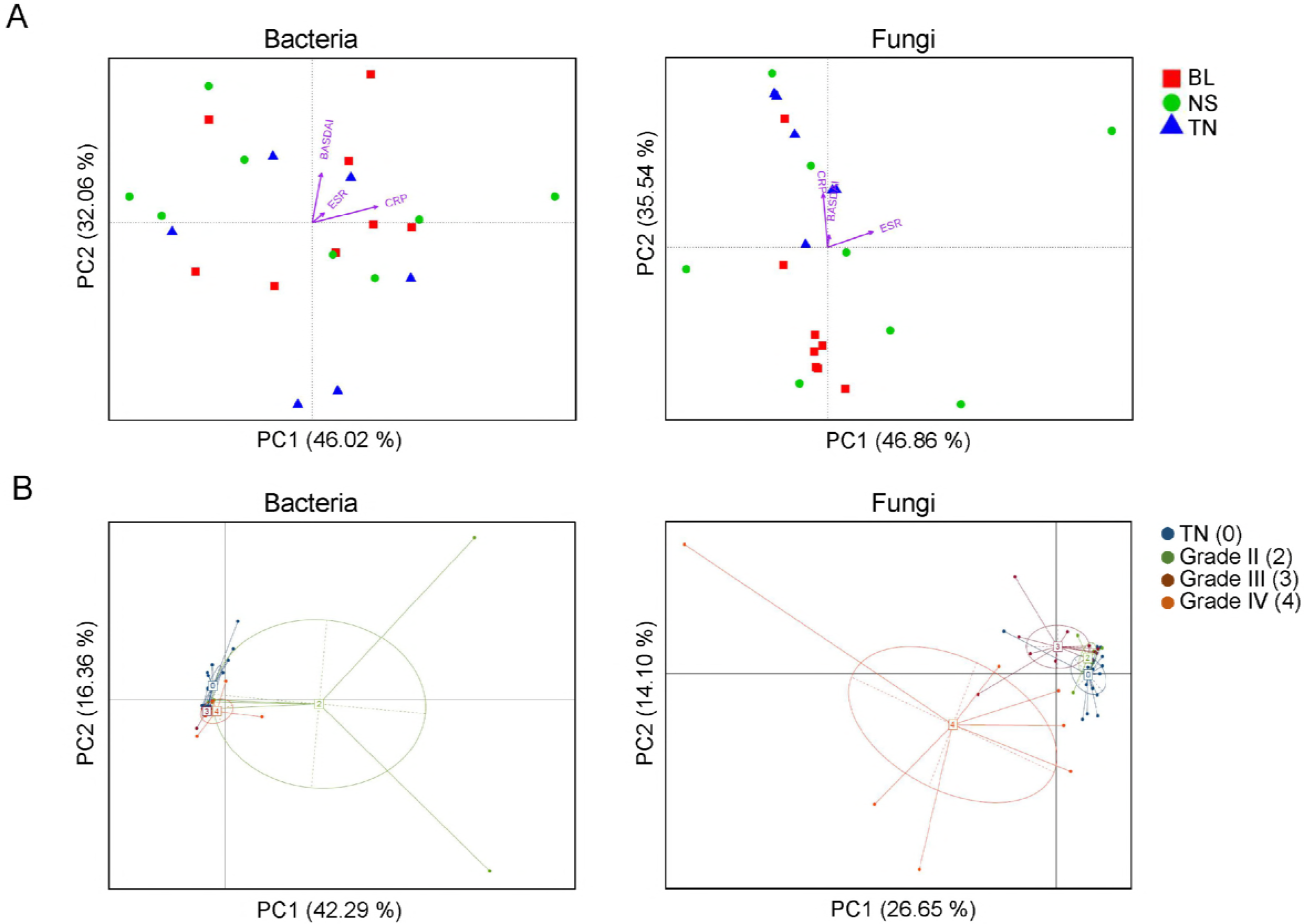
Altered mycobiota in AS patients is associated with disease activities and levels of radiographic damage. (A) The canonical correspondence analysis (CCA) establish the relationship between the disease activity measures and the bacterial and fungal community in AS patients. The direction of arrows indicates correlation with the first two canonical axes and the length of arrows represents the strength of the correlations. (B) PCA of the gut bacterial and fungal genera of AS patients according to their stages of radiographic changes.

## DISCUSSION

In this study, we explored a distinct mycobiota pattern and altered bacteria-fungi interactions in gut of AS patients, which represents a novel research viewpoint of the gut microbiome dysbiosis in AS.

Our finding provided a further confirmation of the alterations in gut microbial groups that might be associated with the development of AS. At the phylum level, increased abundance of *Proteobacteria* and decreased *Bacteroidetes* were found in AS patients, which was proved by previous study in the terminal ileum biopsy specimens of AS patients (14). In addition, there was a greater abundance of *Firmicutes*, *Actinobacteria* and a lower abundance of *Fusobacteria* detected in gut of AS patients, which highly agreed with the results of Wen *et al*.(15). Notably, the decrease of *Bacteroidales* in AS patients was mainly contributed by the depletion of *Prevotella* spp. This result apparently disagreed with Wen *et al*.’s study, in which an increase in the abundance of *Prevotella* spp. was observed in AS patients, although Costello *et al*.’s study supported that the family of *Prevotellaceae* in AS was decreased. *Prevotella* spp. was found with inflammatory properties, as demonstrated by augmented release of inflammatory mediators from immune cells and various stromal cells (28), which suggested that some *Prevotella* strains might be clinically important pathobionts and could participate in human diseases by promoting chronic inflammation. However, in our viewpoint, a depletion of immune-stimulating bacteria in gut may be closely associated with immunodeficiency in human, as supported that *Prevotella* abundance was reduced within the lung microbiota in patients with asthma and chronic obstructive pulmonary disease (29).

Prompted by recent studies in IBD patients (20,31), we profiled the fungal microbiome of the AS patients by sequencing analysis of the ITS2 marker gene, which provided greater resolution of the mycobiome membership compared to metagenomic and 18S rRNA gene sequencing data (20). Interestingly, a more pronounced fungal dysbiosis than bacterial dysbiosis in AS patients was detected in this study. We observed a significant decrease in the diversity of intestinal fungi in these patients. What’s more, the abundance of *Ascomycota* and *Basidiomycota* were strongly negatively correlated with each other and were among the most important discriminative features between AS and HC mycobiota. These results highly agreed with findings in IBD patients, in which the *Basidiomycota*-to-*Ascomycota* abundance ratio differed between patients with IBD and HC (19), suggesting that this imbalance may be either driven by inflammation or involved in the inflammatory process.

Fungi and bacteria coexist in human and animal gut and interact with each other (32-34). Expansion or reduction of fungi can be observed in mice post antibiotics treatment or following antibiotic cessation (35), suggesting a balance between fungal and bacterial microbiota. Our observation of the alterations in the fungi-bacteria diversity balance in AS suggested a modified interkingdom interaction. In addition to differences in the ITS2/16S biodiversity ratio, we noted a disease-specific pattern for the interkingdom network by the spearman correlation analysis. In AS, especially the treatment naïve patients, the number and the intensity of the correlations between fungi and bacteria were increased. The altered biodiversity in bacteria and fungi is associated with new interkingdom interactions that may be involved in the inflammatory process (19). Notably, this interaction in AS patients receiving NB or BL differed significantly from that of the HC and TN groups. Especially in patients treated with BL, the stronger correlations between fungi and bacteria suggested a profound effect of immunosuppressive regimens. Given the limited number of study cases, further large scale studies on the characterization of gut microbiome and mycobiome in AS patients with different therapeutic regimens are necessary.

CRP is well established as biomarker that directly reflect inflammation as acute phase reactants in AS (36). AS patients showed significant correlation between CRP with clinical parameters such as pain, morning stiffness, enthesitis-related local discomfort, BASDAI, BASFI (Bath Ankylosing Spondylitis Functional Index) and BASMI (Bath Ankylosing Spondylitis Metrology Index). In our study, we found a strong positive correlation between serum CRP levels and fungal microbiota in the new cases of AS (TN group), and this pattern was confirmed by the CCA of BASDAI. In contrast, treatment of BL or NS have profound effects on changing of specific gut microbial and fungal groups, which may associate with altered disease activities in AS patients. In addition, it was confirmed that disease activity contributes longitudinally to radiographic progression in the spine of AS patients (37). The structural damage in the spine was found to be associated with the acute phase reactants (APR) CRP and ESR (38-41). We therefore analyzed the gut microbial and fungal microbiota structures according to the radiographs that was scored according to the New York criteria. Interestingly, the gut fungal microbiota of AS patients clustered clearly into three groups, and it was highly correlated with the radiographic assessment. The patients with level III and IV damage in their spines had distinguished fungal microbiota structure when compared with level II or healthy controls, while no significant clustering was observed between the latter two groups. These results suggested an important role of mycobiome in the development of AS.

### Concllusion

In conclusion, our study identified a distinct mycobiota dysbiosis in AS in addition to the alterations in bacterial microbiota. Moreover, we unraveled disease-specific interkingdom network alterations in AS, suggesting that fungi, or the interkingdom interactions between bacteria and fungi, may play a more essential role in AS development. Finally, although our study was not statistically sufficient, we identified some trends suggesting that different therapeutic regimens may induce changing of both bacterial and fungal microbiota in AS.

## MATERIALS AND METHODS

### Study subjects and sample collection

The recruitment of participants and the process of sample collection were depicted in figure S1. Fourty one patients (aged 15 – 58 years) were ultimately recruited from Dalian Municipal Central Hospital and the Second Affiliated Hospital of Dalian Medical University, Dalian, China, from May to September 2017. The disease activity measures of AS patients included the Bath AS Disease Activity Index (BASDAI), AS Disease Activity Index (ASDAS)-C-reactive protein (CRP), CRP, erythrocyte sedimentation rate (ESR), patient’s global assessment and spinal pain (37). And two readers independently scored the radiographs according to the New York criteria, which describes 5 grades of sacroiliitis ranging from 0 to 4 (42).

The fecal samples were collected in Stool Collection Tubes, which were pre-filled with Stool DNA Stabilizer for collection (Stratec, Germany), then frozen and stored at-80 ºC for further use. All subjects were examined clinically before sampling and were subsequently divided into four groups according to different pharmacological therapies: treatment naïve (TN, n=8), patients receiving non-steroidal anti-inflammatory drug (NSAID, n=18) and patients receiving biologics (BL, n=15). The samples of the healthy controls (HC, n=30) were collected during routine physical examination at the Liaoning International Travel Health Care Center, Dalian, China.

The participants with the following diseases were excluded: cardiovascular disease, diabetes mellitus, liver cirrhosis, infections with known active bacteria, fungi, or virus. Those who abused drug or alcohol in the last year, or used antibiotics, probiotics, prebiotics or synbiotics in the month before collection of the fecal samples were also excluded.

### DNA isolation and library construction

The metagenomic DNA in the fecal samples was extracted by the QIAamp DNA stool mini kit (Qiagen, Germany). The purity and concentration of the metagenomic DNA were measured by NanoDrop 2000 spetrophotometer (Thermo, USA).

The V3-V4 region of 16S rDNA (representing bacteria) and the internal transcribed spacer regions 2 (ITS2, representing fungi)^20^ were amplified with the primers (16S: F341 and R806, PCR product: 425 bp; ITS2: ITS3 and ITS4, PCR product: 320 bp). Primer sets were modified with Illumina adapter regions for sequencing on the Illumina GAIIx platform, and reverse primers were modified with an 8-bp Hamming error-correcting barcode to distinguish among samples. The DNA template (100 ng) was combined with 5 μL PCR buffer, 1 μL dNTPs, 0.25 μL HotStarTaq^®^ Plus DNA Polymerase (Qiagen), and 2.5 pmol of each primer in 50 μL total volume. Reactions consisted of an initial step at 95 °C for 5 min; 25 (16S rDNA) or 38 (ITS2 rDNA) cycles of 94 °C for 45 s, 55 °C for 45 s and 72 °C for 60 s; and a final extension at 72 °C for 10 min. DNA products were checked by 1.5% (w/v) agarose gel electrophoresis in 0.5 mg/mL ethidium bromide and purified with the Qiaquick gel extraction kit (Qiagen).

### Bioinformatics analysis

Sequences of the V3-V4 region of 16S rDNA and ITS2 were detected using an Illumina HiSeq PE250 platform (reconstructed cDNA sequence: 2 × 250 bp, Novogene Bioinformatics Technology Co. Ltd, Beijing). Ribosomal Database Project (RDP) Classifier 2.8 was used for taxonomical assignment of all sequences at 50% confidence after the raw sequences were identified by their unique barcodes. OTUs present in 50% or more of the fecal samples were identified as core OTUs. PLS-DA of core OTUs was performed using Simca-P version 12 (Umetrics), and a heat map was generated with Multi-Experiment Viewer (MeV) software to visualize and cluster the fungal community into different groups. Community diversity was measured by the Shannon-Weiner biodiversity index (Shannon index).

### Statistical analysis

All data were evaluated as mean ± SEM. Statistical analysis of the quantitative multiple group comparisons was performed using one-way analysis of variance (and non-parametric), followed by wilcox’s test; when two groups were compared, the non-parametric *t*-test was performed with the assistance of GraphPad Prism 6 (Graph Pad Software, La Jolla, CA, USA). Results were considered to be statistically significant with p < 0.05. * p < 0.05; **p < 0.01; *** p < 0.001.

### Ethics statement

This study protocol was approved by the Ethics Committees of all participating hospitals including Dalian Municipal Central Hospital and the Second Affiliated Hospital of Dalian Medical University, Dalian, China. All the procedures were performed in accordance with the guidelines approved by the Ethics Committee of Dalian Medical University, China. After receiving a written description of the aim of this study, all participants gave written informed consent prior to enrollment.

## ACKNOWLEDGMENTS

This research was funded by the National Natural Science Foundation of China (No. 81671606), the program of Liaoning Distinguished Professor (Liao taught 2018-2020), the China Postdoctoral Science Foundation (2016M601317, 2018T110225), and the Research Foundation from the Department of Education, Liaoning Province, China (L2016003). This work was supported by Liaoning Provincial Program for Top Discipline of Basic Medical Sciences.

---

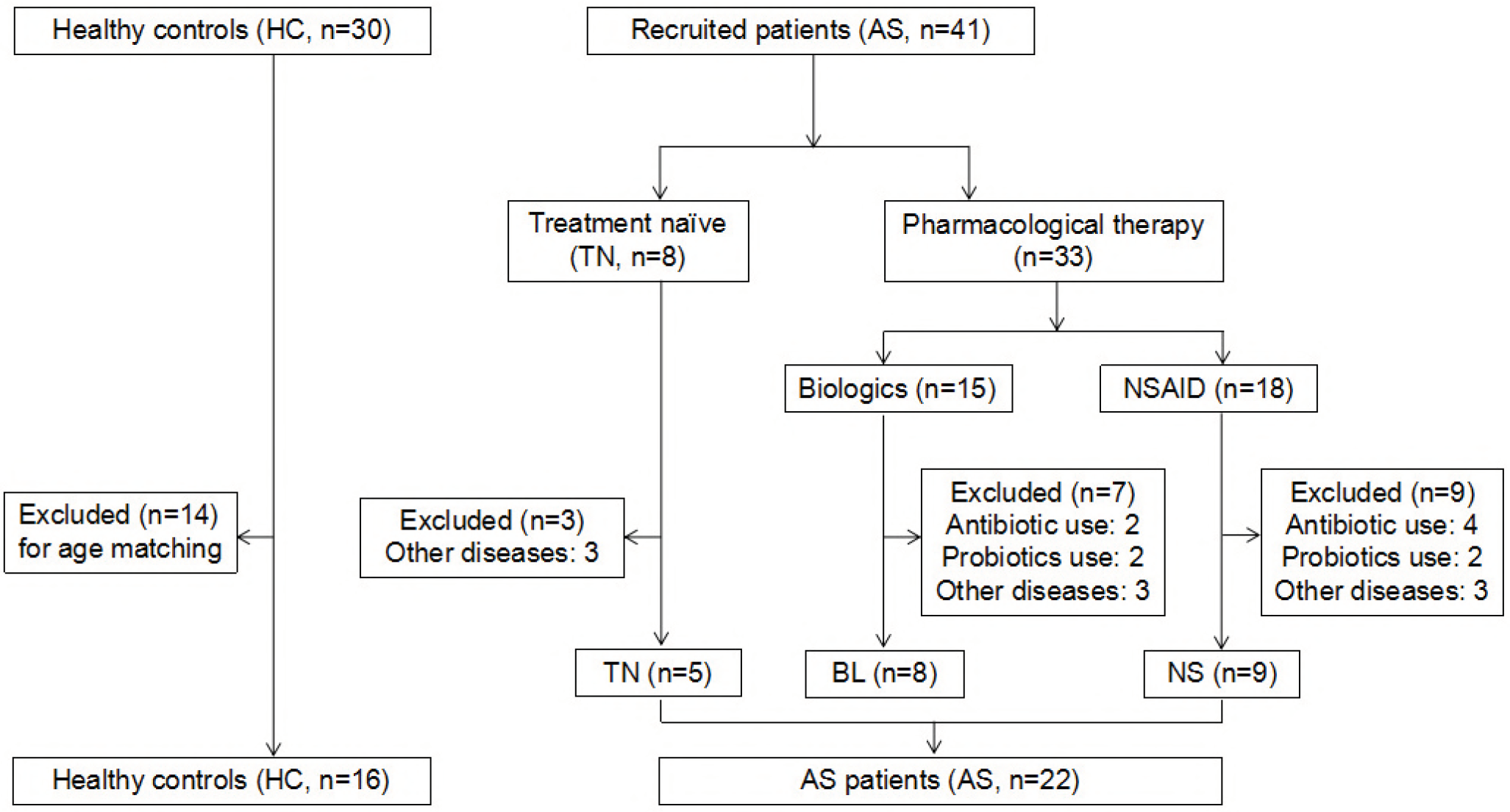

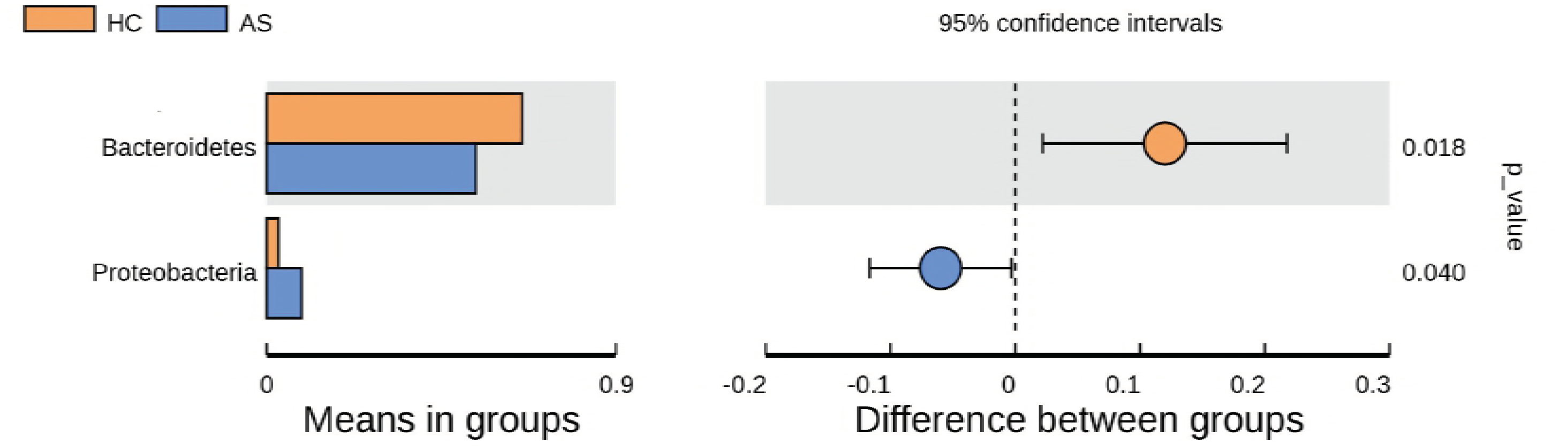

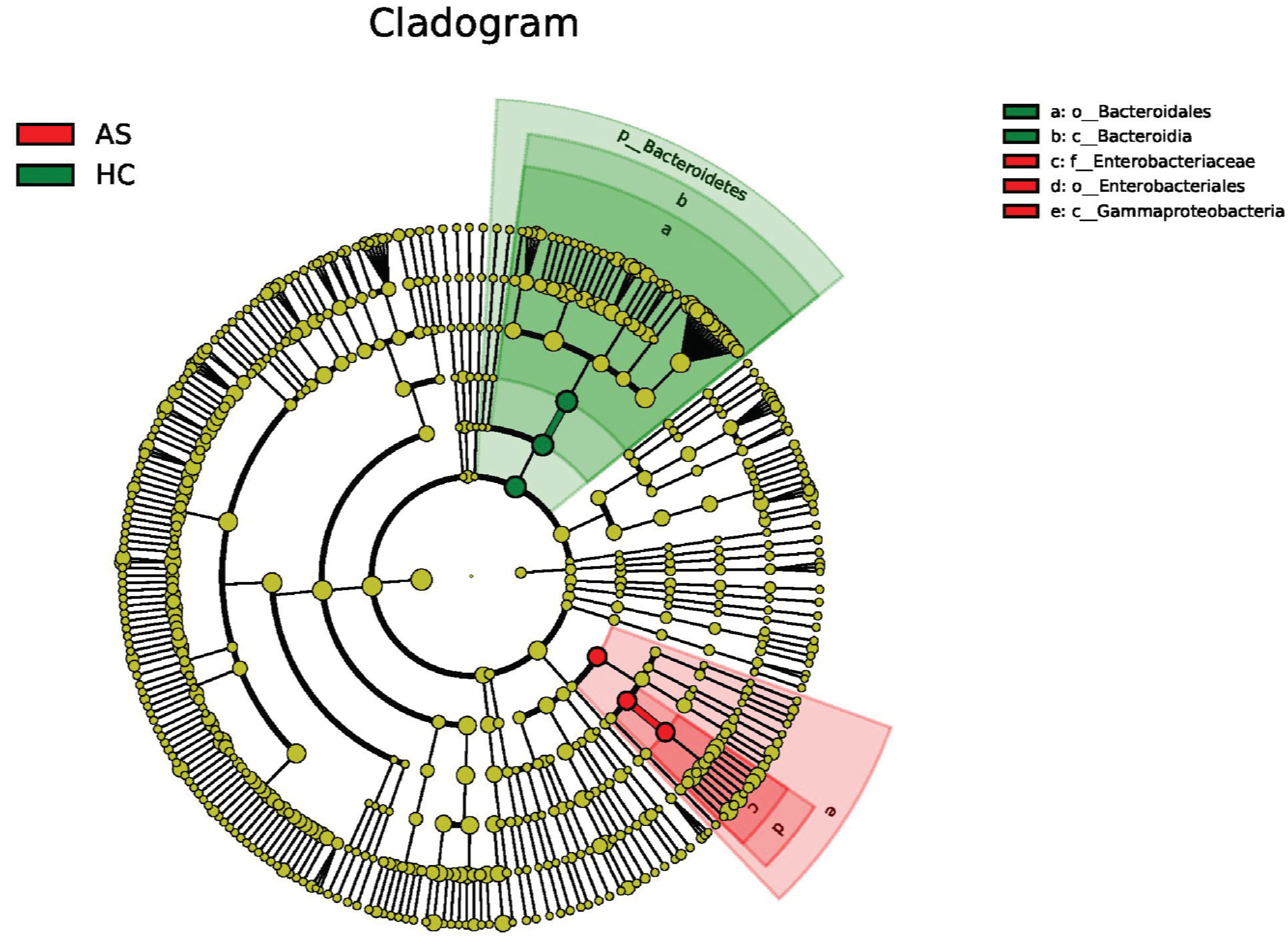

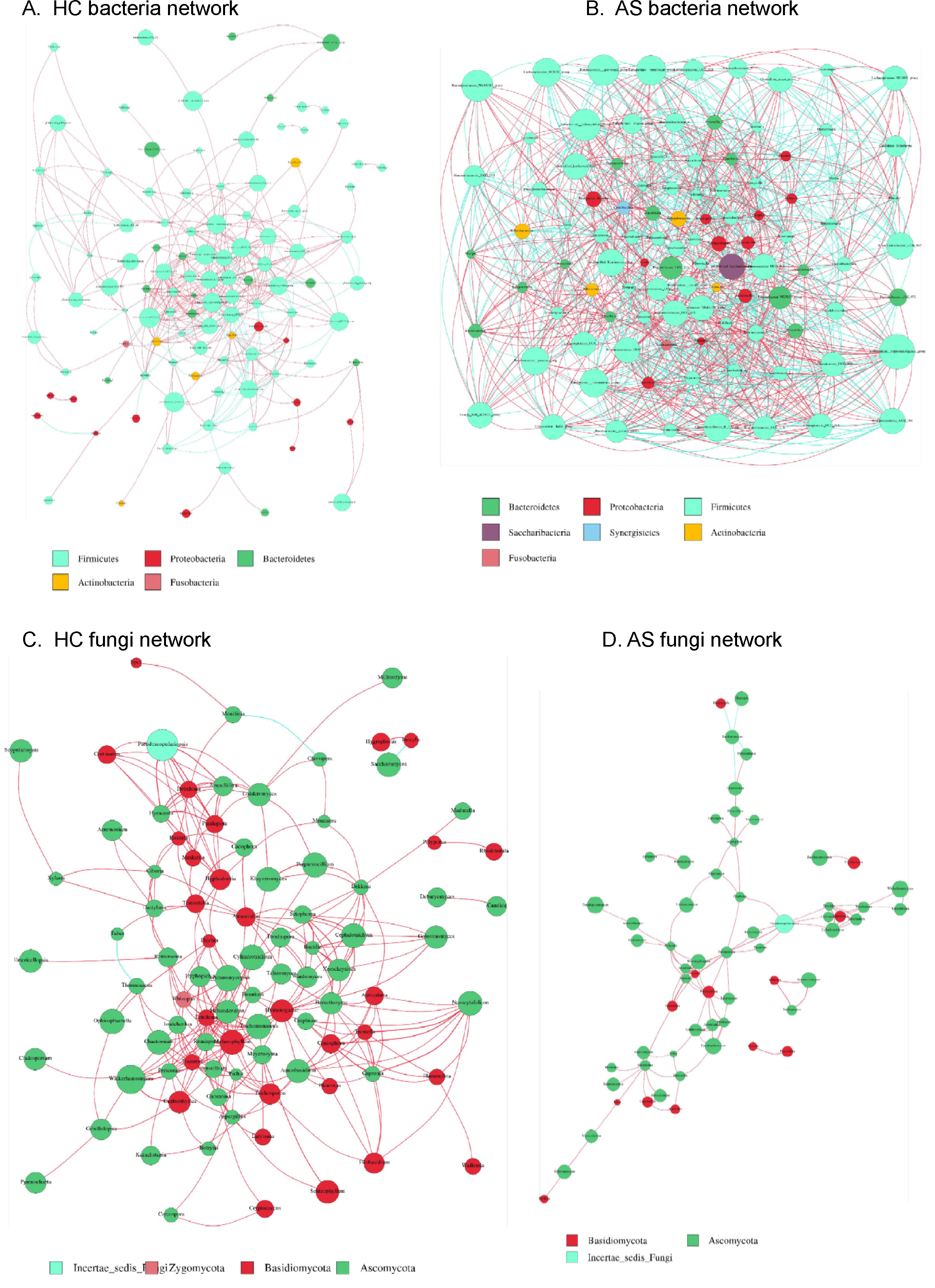

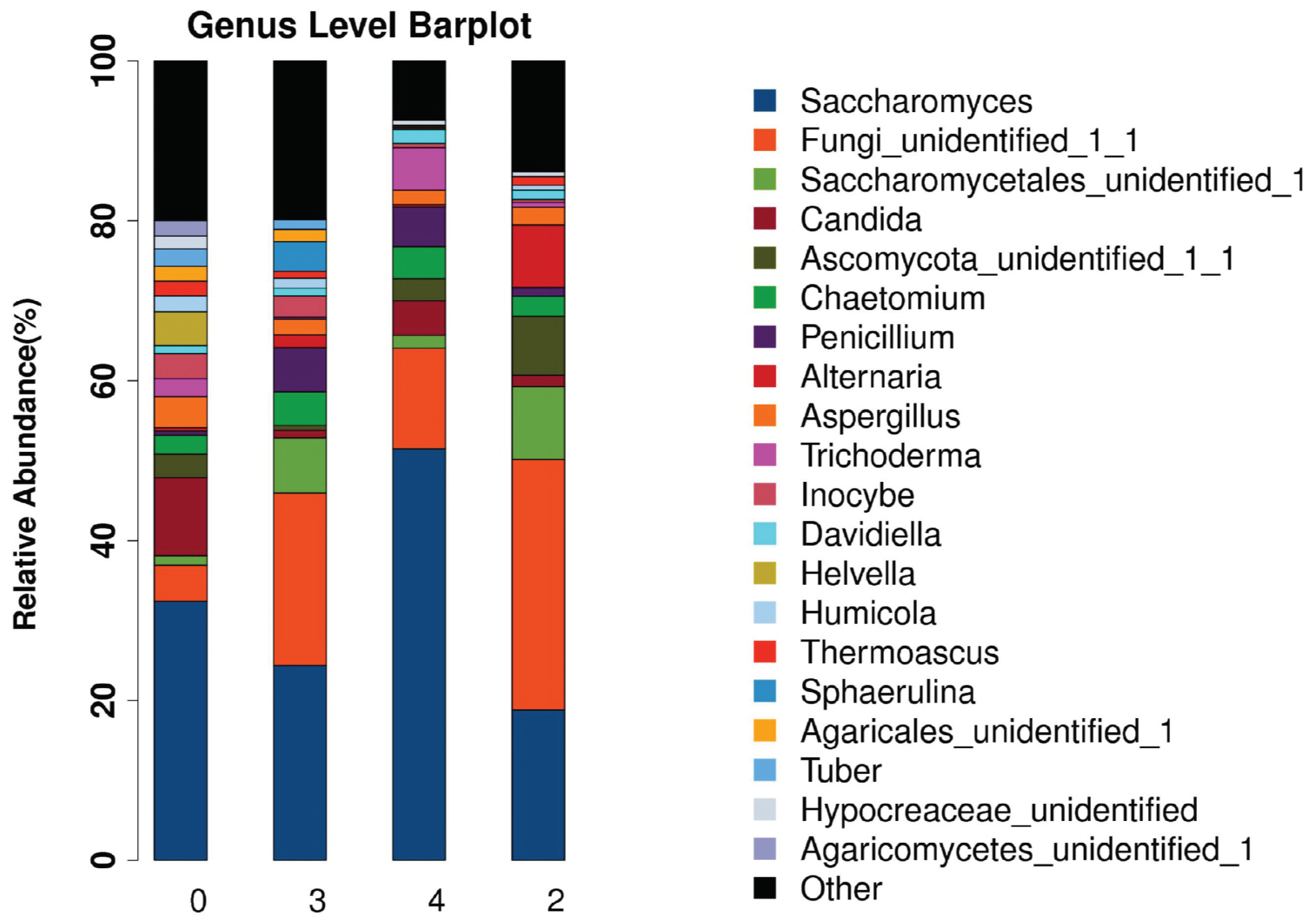

